# DIET IMPACTS THE EFFECTS OF THERMAL STRESS ON INFECTION OUTCOMES

**DOI:** 10.1101/2025.10.26.684613

**Authors:** Biswajit Shit, Selah Makinishi, Tanmay Singh, Imroze Khan

**Affiliations:** Trivedi School of Biosciences, Ashoka University, Sonepat, Haryana, India; Centre for Climate Change and Sustainability, Ashoka University, Sonepat, Haryana, India

**Author notes:** Equal contribution (Selah Makinishi, Tanmay Singh).

**Keywords:** Dietary Manipulation, Infection, Fertility, Starvation, Mating status

## Abstract

**Background:** Climate change can lead to warmer environments, facilitating higher pathogen growth and increasing disease burden. Climate change can also reduce the nutritional value of food resources, causing malnutrition, which can lead to impaired immunity and disease vulnerability. However, despite these associations, no experiments have tested how dietary manipulations and temperature variations interact to drive the effects of pathogenic infections. In this work, we conducted laboratory experiments with the model insect *Drosophila melanogaster* to test these effects.

**Results:** Virgin flies raised on low-carbohydrate diets exhibited improved post-infection survival rates and more resilience to pathogen burden compared to those reared on standard or high-carbohydrate diets. However, such a survival advantage of the low-carbohydrate diet was lost in mated individuals. Notably, temperature stress decreased survival rates in flies only when they were reared on a standard diet. This effect was not observed in flies raised on high- or low-carbohydrate diets, indicating that the impact of temperature on infection responses might depend on diet. Besides, flies developing on a low-carbohydrate diet showed greater fertility at ambient temperatures than those on other diets, indicating an apparent lack of fitness costs associated with improved post-infection response. Also, high-carbohydrate diets conferred greater resistance to starvation, suggesting differences in resource allocation among somatic maintenance tasks that could increase vulnerability to pathogens.

**Conclusion:** Overall, these findings imply a complex interaction between diet, temperature, and fitness traits that influences variation in infection responses. As many organisms around the world face environmental warming and poor diets, examining these patterns and processes can provide a deeper understanding of variations in health and disease responses amid a changing climate.

## INTRODUCTION

Climate change may heighten the prevalence and severity of infectious diseases, significantly impacting global health and disease management (Altizer *et al*., 2013). Two primary factors likely driving these climate change effects are the warming of the environment and diet quality (Wu *et al*., 2016). Previous research indicates that elevated temperatures can increase vulnerability to pathogenic infections by interfering with various immune components (Schade *et al*., 2014), as exemplified by reduced antimicrobial peptide (AMP) expression in mosquitoes (Murdock *et al*., 2012) and fruit flies (Linder *et al*., 2008), diminished antibody production in turtles (Dang *et al*., 2015), to theoretical predictions of weakened antifungal immunity in frogs (Raffel *et al*., 2013) and human innate immune responses (Presbitero *et al*., 2021). Additionally, global warming can facilitate the spread of vector-borne, water-borne, air-borne, and food-borne pathogens, resulting in numerous human diseases, including Malaria, Dengue, Typhoid fever, and Echinococcosis, among others (Mora *et al*., 2022).

The fate of pathogenic infections can also significantly hinge on the quality of the diet. As infections and the resulting immune responses elevate metabolic rates, the energy required to combat pathogens is likely to be demanding, especially when nutrition is inadequate (Galenza & Foley, 2019). Additionally, variations in macronutrient intake can greatly influence post-infection responses in organisms (Graham *et al*., 2014; Kay *et al*., 2014; Tsai *et al*., 2022). For example, consuming a high-protein diet might heighten susceptibility to pathogens by reducing the expression of immune-related genes (Dinh *et al*., 2019; Ponton *et al*., 2020). However, dietary regulation is also a key determinant of thermal tolerance, as it contributes to maintaining metabolic homeostasis in an organism under thermal stress. Indeed, previous studies have suggested that protein restriction can impair thermal tolerance (Strilbytska *et al*., 2021) and elevates metabolic rates at higher temperatures, increasing the overall energy requirements of organisms (Alton *et al*., 2020). In fact, a protein-rich diet provides improved resistance to heat knockdown in female fruit flies (Andersen *et al*., 2010). Given such contrasting effects of dietary protein content during thermal stress versus infection stress, it remains unclear how organisms respond when they face pathogenic infection and elevated temperatures simultaneously. Understanding the interplay among diet, temperature, and immunity is thus essential to predict infection outcomes in a changing climate.

In this work, we have explored how dietary manipulations by altering the macronutrient composition (e.g., carbohydrate vs protein, henceforth C:P ratios) drive the infection outcome of the host under elevated temperatures. Since mated individuals can differ from virgin individuals in post-infection outcomes due to a resource allocation trade-off between immunity and reproduction (Schwenke *et al*., 2016), we compared the effects of infection as a function of the mating status. To test if resource allocation trade-offs between various fitness traits explain the variation in the observed infection outcomes, we also checked fertility and starvation resistance across dietary treatments and temperature conditions. Overall, our findings indicate that the effects of temperature on pathogen clearance capacity and post-infection survival differ significantly depending on dietary composition and mating status, exhibiting complex relationships with alterations in reproductive ability and body condition.

## MATERIALS AND METHODS

### Post-infection survival and bacterial load quantification

We used an outbred population of *Drosophila melanogaster* for our experiments. This population is maintained on a standard cornmeal-sugar-yeast diet (Mudunuri *et al*., 2024) at a constant temperature of 25°C on a 24-hour light cycle at 60% humidity. Additionally, we used only females for our study since previous studies have suggested that females generally invest more in immunity to maximize their breeding time (Belmonte *et al*., 2020). Briefly, for the experiments, we reared and assayed the females in 3 dietary treatments with altered carbohydrate vs protein ratio (henceforth, C:P), such as HighC (C:P = 8:1), Standard (C:P = 4:1), and lowC (C:P = 1:1) (Ponton *et al*., 2020), and 2 temperature conditions, namely ambient (25°C) and warm (29°C). We used the gram-negative bacterial pathogen *Providencia rettgeri*, a natural pathogen for *Drosophila melanogaster* that causes pronounced pathogenicity (Galac & Lazzaro, 2008). We infected 3 days old adult virgin flies, using the septic injury method by pricking the thorax region with a 0.1 mm minutien pin (Fine Science Tools, Canada) dipped into a bacterial suspension (OD600= 20; equivalent to ∼250 bacterial cells/fly) (Shit *et al*., 2022). We infected 75 flies for each treatment and distributed them in 5 replicate food vials, each containing 15 females. Sham infections were also carried out by pricking with sterile 1X PBS as a procedural control (n= 75 flies/treatment; each treatment had five replicate vials with 15 females. To generate the mated flies, we paired 2-day-old virgin males and females in a single vial for each treatment (15 vials/treatment) and allowed them to mate for 24 hours with a mating pair of 5 individuals in each vial. Following this, on the next day, we separated all the females from each of the vials and pooled them together to perform live infection and sham infection using the previously described methods and sample sizes. To check the host’s ability to suppress bacterial growth under different diet-temperature combinations, we isolated 3 females for each replicate vial across treatments to quantify their bacterial load at 20 hours post-infection, where mortality typically begins in flies infected with *P. rettgeri*. First, we surface sterilised individual flies and homogenised them, followed by serial dilution of the fly homogenate up to 10^5-fold. 4μL of the aliquot from each of the dilutions was added to LB agar plates, and the plates were kept for overnight incubation at 37°C to count the bacterial colonies as described by Shit et al. (2022) (Shit *et al*., 2022). The method was same in both virgin and mated females.

We recorded the post-infection survival of the remaining flies by counting the number of deaths every 4 hours for the next 5 days (n= 60 flies/treatment; each treatment had five replicate vials with 12 females). We also collected the post-infection data for mated individuals using the same procedure. We analysed fly post-infection survival as a function of dietary treatment and temperature using the Cox proportional hazard model, implementing the R package ‘survival’ (Therneau & Grambsch, 2000), specified as ‘Post-infection survival ∼ Dietary treatment× temperature’. We analysed the virgin and mated flies separately, as they were assayed on two different days. We analysed the bacterial load data using a generalised linear model fitted to a gamma distribution (lowest AIC compared to the other relevant distribution; AIC =365.21 (for virgin individuals), AIC = 406.79 (for mated individuals)) using R package ‘lme4’, with ‘diet’ and ‘temperature’ as fixed effects (model: CFU∼ Dietary treatment× Temperature; family = Gamma).

### Measurement of fertility and body condition

To check whether variation in infection response as a function of diet and temperature is also associated with the changes in reproductive capacity and body condition, we estimated the number of offspring produced and lifespan under starvation (Rion & Kawecki, 2007)of the females from each treatment. To measure reproductive output, we paired 3-day-old virgin 5 females and 5 males in a single vial for each treatment (n=7 vials/treatment) and allowed them to mate for 24 hours. Following this, we separated 5 females from each vial under C02 anaesthesia and distributed them individually to oviposit. We discarded the females after 24 hours and kept the vials with eggs to allow complete eclosion. We counted the number of adult offspring from each vial on next day post-oviposition. We analysed the fertility data using a generalised linear model with a ‘negative binomial distribution’ since the data was not normally distributed and ‘negative binomial’ had the least AIC value and lowest overdispersion index (AIC value = 1500.1, overdispersion index = 0.59). For this analysis, we used R package ‘lme4’, with ‘diet’ and ‘temperature’ as fixed effects (model: Fertility∼ Dietary treatment × Temperature; family = Negative binomial).

To estimate starvation resistance, we housed 3-day-old virgin adult females in vials containing 2 ml 1.2% non-nutritive agar gel at a density of 8 females per vial (n=10 vials/treatment) (Singh *et al*., 2025). The presence of agar gel in the vials ensured that the flies had ad libitum access to water during the starvation assay. We monitored the vials every 4 hours to record the number of dead flies until the last fly perished. Surviving flies were transferred to fresh agar vials every 48 hours. We analysed the starvation resistance data using the Cox proportional hazard model, taking ‘Dietary treatment’ and ‘Temperature’ as fixed effects (the specified model was survival ∼ Dietary treatment × Temperature).

We used statistical software R (v4.3.1, R core team 2022) for all the analyses.

## RESULTS

### Rearing in low-carbohydrate media leads to increased post-infection survival

At ambient temperature, virgin flies fed a low-carbohydrate diet exhibited the highest survival rate post-infection (∼75%) (**Fig**. 2A, **Table** S1). In contrast, those raised on high-carbohydrate diets experienced the lowest survival rates against *P. rettgeri*. Virgin flies reared on a low-carbohydrate diet showed the highest post-infection survival rate under warmer condition*s* as well. However, unlike ambient conditions, individuals reared on the standard diet had the lowest survival (**Fig**. 2A, **Table** S1). Despite the changes in survival patterns, there were no differences in bacterial loads across diet and temperature treatments, suggesting that the observed differences in post-infection survival may reside in their divergent ability to withstand the bacterial load (**Fig**. 2B, **Table** S2).

**Figure 1:**
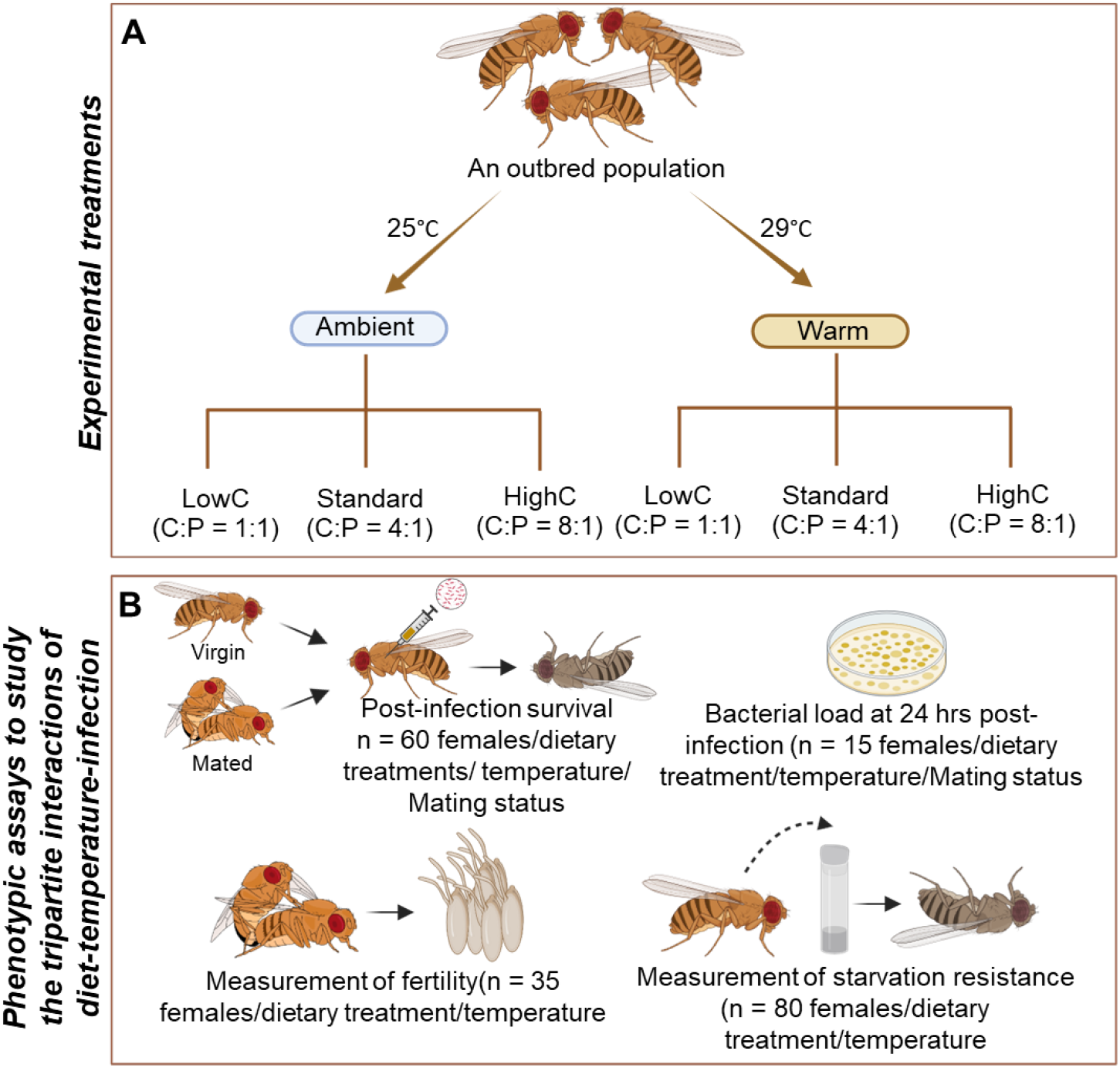
Outline of the experimental design to study the impact of tripartite interactions between diet, temperature, and mating status in response to pathogens: (A) Generation of the experimental female flies across treatments for all the assays. Briefly, the flies from an outbred population are either reared under ambient temperatures (25°C) or warm temperatures (29°C) and three dietary conditions-Low-carbohydrate (C): Protein (P) ratio (LowC), Standard diet, and High-C:P ratio (HighC) (6 different treatments in total); (B) Different phenotypic assays to know the effect of interactions between dietary treatments and temperature—e.g., response to pathogenic infections and bacterial load in virgin vs mated females, female fertility, and starvation resistance of virgin females. The replicate size of each experiment has been provided in panel B.

**Figure 2:**
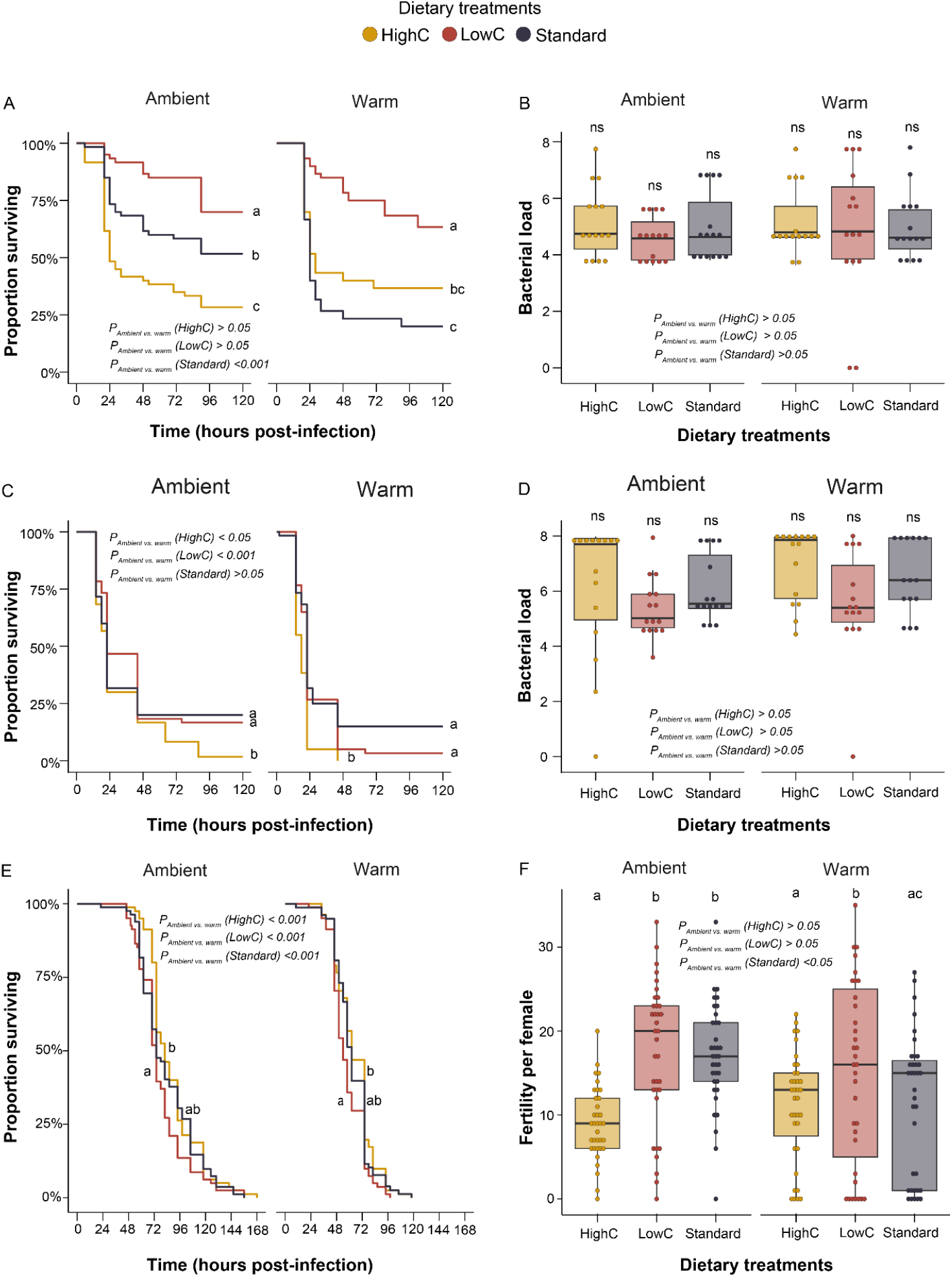
(A) Survival curves of female flies from different dietary treatments (Low-carbohydrate (LowC), High-carbohydrate (HighC) and standard (Standard) diets) and different temperature conditions (Ambient and Warm) for (A) virgin flies (C) mated flies following infection with *P. rettgeri* (n = 60 females/dietary treatment/temperature/mating status). (B) & (D) Bacterial loads after 20 hours of *P*.*rettgeri* infection for virgin & mated flies (n = 15 females/dietary treatment/temperature/mating status). (E) starvation resistance of female flies from different dietary and temperature treatments mentioned for the post-infection survival assay (n = 80 females/dietary treatment/temperature). (F) Fertility of the females from different dietary and temperature treatments (n = 35 females/dietary treatment/temperature). In panels B & D, each data point represents the whole-body bacterial load of a single fly. In all the panels, significantly different dietary treatments (P< 0.05) are indicated by distinct alphabets, and ‘ns’ indicates that the comparisons are not significantly different. Significantly different temperatures are marked by the P-values mentioned in the panels. For panels A, C & E, the data are analysed using the Cox proportional hazard model. For panels B & D, the data are analysed using a generalised linear model fitted to a gamma distribution. For panel F, the data are analysed using a generalised linear model fitted to a negative binomial distribution.

Furthermore, we could detect the effect of temperature only on flies reared under standard conditions, where they became more vulnerable to pathogenic infections at warmer temperatures. Surprisingly, there was no effect of temperature on infection susceptibility for either the high- or low-carbohydrate groups, indicating that the impact of temperature can be diet-specific (**Table** S1).

By and large, mating increased post-infection susceptibility, and none of the dietary manipulations showed an effect of temperature on infection susceptibility (**Fig**. 2A-B, **Table** S1). No survival benefit was observed in the case of low-carbohydrate food, as seen earlier in the case of virgin flies (**Fig**. 2C, **Table** S1). However, high-carbohydrate treatment again resulted in the lowest post-infection survival rates, regardless of the temperature treatments (**Fig**. 2A-B, **Table** S1). Mated females harboured greater pathogen loads than virgins (**Fig**. 2D, **Table** S2); but there was no effect of temperature or diet on bacterial load (**Fig**. 2D, **Table** S2).

### Developing under high-carbohydrate media increases starvation resistance

We wanted to check the female’s response to starvation after rearing under different dietary manipulations and temperature variations. Our results suggested that individuals reared under high-carbohydrate food had significantly higher starvation resistance compared to the low or standard carbohydrate food when they were grown in ambient conditions (**Fig**. 2E, **Table** S3). The result was similar when the individuals were developed in warm temperatures (**Fig**. 2E, **Table** S3). Interestingly, ambient-temperature reared individuals had better starvation resistance compared to the warm temperature-reared individuals for flies developed under all three dietary treatments (**Fig**. 2E, **Table** S3)

### Developing under low-carbohydrate media increases fertility

Next, we wanted to check the female’s ability to produce offspring or the fertility after rearing under different dietary manipulations and temperature variations. Our results suggested that the individuals reared under a high-carbohydrate diet had significantly lower fertility than the low-carbohydrate and standard diets, and there was no difference between the low-carbohydrate and standard diets (**Fig**. 2F, Table S4). A similar result was observed in warm-reared individuals (**Fig**. 2F, Table S4), where individuals reared on a low-carbohydrate diet had higher fertility compared to their high-carbohydrate counterparts, although the trend was not statistically significant. Interestingly, there was no effect of temperature on the fertility of individuals developed under low or high-carbohydrate food, but standard food-reared individuals had lower fertility when they developed in warm temperatures (**Fig**. 2F, Table S4) compared to their rearing under ambient temperature. Overall, our results indicated that the low-carbohydrate diet was beneficial for increasing reproductive fitness in both ambient and warm temperature conditions.

## Discussion

In this study, we examined the interactions between dietary manipulation and temperature variation on the organism’s ability to fight against *P*.*rettgeri* infection, followed by estimating changes in fitness traits such as fertility and stress resistance to explain the variation in infection outcomes. Specifically, we found that virgin flies reared on a low-carbohydrate diet showed improved post-infection survival and infection tolerance after *P. rettgeri* infection compared to those raised on standard or high-carbohydrate diets, but the mated individuals lost the post-infection survival benefit of a low-carbohydrate diet. Also, rearing under a standard diet reduces post-infection survival when virgin individuals are exposed to higher temperatures, but this does not apply to mated flies or those reared under low- and high-carbohydrate diets. Diet manipulations and reproductive status thus critically influenced the effects of thermal stress on infection outcomes. Interestingly, in none of the treatments did pathogen burden predict pathogen vulnerability—i.e., we did not observe any changes in bacterial load due to diet and temperature manipulations, despite differences in post-infection survival rates. This suggests the need for further research to determine whether the effects of diet, temperature, and life history primarily operate by altering pathogen tolerance rather than by actively reducing the pathogen load through resistance mechanisms to prevent immunopathological effects (Seal *et al*., 2021; Prakash *et al*., 2025).

Our results were consistent with one of the previous findings, which showed high-glucose diets to have adverse effects on the quality of immune defence against the bacteria *P. rettgeri* (Fellous and Lazzaro, 2010). A high-carbohydrate environment can reduce the constitutive expression of antimicrobial peptides (AMPs) like IMD-pathway responsive Diptericin, which are critical for survival against *P. rettgeri* (Fellous and Lazzaro, 2010; Shit et al., 2022). By contrast, flies maintained on a low-carbohydrate diet perhaps had more opportunity to access protein, which may translate into more AMP production (Darby *et al*., 2024), thereby rendering the flies more resistant to *P. rettgeri*. Indeed, previous studies have demonstrated that fruit flies raised on a low-carbohydrate diet exhibited increased Drosocin expression when infected with P. rettgeri (Darby *et al*., 2024), which is recognised as one of the IMD-responsive AMPs that protects against *P. rettgeri* in females (Shit et al. 2022). Parallely, a high-carbohydrate diet may also induce insulin resistance in flies, which has been previously linked with reduced hemolymph melanisation and the expression of AMPs (Musselman *et al*., 2018; Darby & Lazzaro, 2023). Flies reared in a high-carbohydrate diet in our study may have encountered a negative feedback loop between insulin signaling and the signaling pathway responsible for innate immunity (Musselman *et al*., 2018), where induced insulin signalling can suppress the transcription factor Forkhead box O (FOXO), leading to reduced expression of several antimicrobial peptides such as Diptericin, Cecropin A1, and Drosomycin, which results in higher pathogen vulnerability (Darby & Lazzaro, 2023).

One of the most striking outcomes of our results is that although the individuals reared under standard food had higher susceptibility against *P. rettgeri* in warm temperatures, the post-infection outcome is similar in both low-carbohydrate and high-carbohydrate reared individuals. In fact, any deviation from the standard diet (i.e., both low- and high-carbohydrate diets) significantly improved post-infection survival at high temperature, thereby mitigating temperature-induced decline in post-infection survival. We speculate that although high temperature depletes body energy reserves in *D. melanogaster* (Klepsatel *et al*., 2019), rearing on a low-carbohydrate diet may help them to regain the temperature-induced loss of post-infection survival by increasing the expression of immune genes (Darby *et al*., 2024). On the other hand, individuals reared under a high-carbohydrate diet might be promoting more energy storage in the fat body by stimulating both carbohydrate and lipid storage pathways (Rajan & Perrimon, 2013; Lee & Jang, 2014). This may help them effectively offset energy depletion during infection (Hall *et al*., 2024) and temperature stress (Klepsatel *et al*., 2019), improving post-infection health at warm temperatures. Moreover, we observed a general decline in post-infection survival with mating, regardless of dietary treatments. This also led to the loss of the survival benefit that we observed in virgin flies reared under a low-carbohydrate diet. This is expected as the mating and subsequent investment in egg production are costly, reducing resources available to mount an effective immune response (Short et al., 2012).

In addition to poor post-infection survival, individuals on a high-carbohydrate diet also experienced reduced fertility compared to those on other dietary regimes, suggesting that the immunity-reproduction trade-off may not explain the variation in infection outcomes. Lower fertility with a high-carbohydrate diet is, in fact, consistent with a previous finding that showed fruit flies reared under a high-glucose diet produced fewer offspring (Howick & Lazzaro, 2014), which may arise from protein deficiency, causing a significant reduction in yolk protein concentration in adult female flies, a factor essential for egg development (Sisodia & Singh, 2012). These individuals also showed better starvation resistance, suggesting better energy storage, and corroborate previous experiments indicating that a low-protein diet enables fruit flies to deposit more lipid and lower the critical body weight to tolerate starvation stress (Lee & Jang, 2014). This may have eventually helped them utilise endogenous fuels more effectively and become more starvation-resistant (Lee & Jang, 2014). Instead of an immunity-reproduction trade-off, our results thus reveal a trade-off among somatic maintenance functions in flies reared under a high-carbohydrate diet (McKean et al., 2008)—such as immunocompetence vs. stress response—that might explain the low post-infection survival rates observed in these flies. However, this hypothesis needs further exploration.

In summary, our results indicate that assessing the impact of climate warming on host-pathogen interactions may not be straightforward, underscoring the need for information on various other health and ecological factors (e.g., reproductive status, resource quality) that shape differences in immunocompetence and vulnerability to pathogens. Since many organisms worldwide are exposed to environmental warming and poor diet, understanding these patterns and processes will offer better insights into variations in health and disease responses in a changing climate.

## Acknowledgement

We thank Saubhik Sarkar, Srijan Seal, and Dipendra Nath Basu for feedback on the manuscript; Sudipta Tung for generously providing us with the fly lines; Sriparna Bhowmick for laboratory assistance; Susheel Dahiya and Darshan Ghelawat for assistance in providing the materials for the experimental setup.

## Author contribution

IK conceived the experiments; IK, BS, TS designed the experiments; BS, TS and SM performed the experiments; BS analysed the data; IK acquired the funding and provided resources and consumables. IK and BS drafted the manuscript with additional input and comments from TS and SM. All authors agreed on the final version of the manuscript.

