## Supplemental information for "DIET IMPACTS THE EFFECTS OF THERMAL STRESS ON INFECTION OUTCOMES"

**SUPPLEMENTARY INFORMATION**

**Table S1**. Summary of Cox proportional hazard model, fitting the model to estimate response to 20 OD (equivalent to ~250 bacterial cells/fly) of gram-negative *P. rettgeri* bacterial infection in virgin and mated flies from an outbred population. The model is specified as survival ~ temperature x dietary treatment, with 'temperature' and 'dietary treatment' as fixed effects. The table shows model output (ANOVA) for survival post-infection in virgin and mated flies. Statistically significant p-values are highlighted in bold.

| | | | | | | |
|---|---|---|---|---|---|---|
| | | ***Tested effect*** | *loglik* | *Chisq* | *Df* | *p value* |
| | *Full model* | *Temperature (T)* | -1094.9 | 4.5498 | 1 | **0.022** |
| | | *Dietary treatment (DT)* | -1071.2 | 47.5251 | 2 | **<0.001** |
| | | *T × DT* | -1064.7 | 13.0451 | 2 | **0.001** |
| | *Post hoc to compare dietary treatments across Temperatures* | *T* | *loglik* | *Chisq* | *Df* | *p value* |
| | | *Ambient* | | | | |
| | | *LowC - HighC* | -259.39 | 27.782 | 1 | <**0.001** |
| | | *LowC - Standard* | -211.68 | 5.7184 | 1 | **0.017** |
| ***Virgin*** | | *HighC - Standard* | -312.70 | 8.8861 | 1 | **0.003** |
| | | *Warm* | | | | |
| | | *LowC - HighC* | -262.57 | 13.229 | 1 | **<0.001** |
| | | *LowC - Standard* | -292.88 | 32.898 | 1 | **<0.001** |
| | | *HighC - Standard* | -367.71 | 3.0444 | 1 | 0.081 |
| | Post hoc to compare temperatures across dietary treatments | *DT* | *loglik* | *Chisq* | *Df* | *p value* |
| | | *LowC* | -183.74 | 0.7903 | 1 | 0.374 |
| | | *Standard* | -328.04 | 16.482 | 1 | **<0.001** |
| | | *HighC* | -350.75 | 0.8615 | 1 | 0.353 |
| | | ***Tested effect*** | loglik | *Chisq* | *Df* | *p value* |
| | *Full model* | *Temperature (T)* | -1669.4 | 9.7810 | 1 | **<0.001** |
| | | *Dietary treatment (DT)* | -1659.0 | 20.7685 | 2 | **<0.001** |
| | | *T × DT* | -1656.9 | 4.2563 | 2 | 0.119 |
| | *Post hoc to compare dietary treatments across Temperatures* | *T* | *loglik* | *Chisq* | *Df* | *p value* |
| | | *Ambient* | | | | |
| | | *LowC - HighC* | -437.79 | 5.0362 | 1 | **0.025** |
| | | *LowC - Standard* | -409.14 | 0.4047 | 1 | 0.525 |
| ***Mated*** | | *HighC - Standard* | -433.94 | 2.638 | 1 | 0.104 |
| | | *Warm* | | | | |
| | | *LowC - HighC* | -450.26 | 13.709 | 1 | **<0.001** |
| | | *LowC - Standard* | -439.65 | 1.324 | 1 | 0.250 |
| | | *HighC - Standard* | -436.30 | 17.415 | 1 | **<0.001** |
| | Post hoc to compare temperatures across dietary treatments | *DT* | *loglik* | *Chisq* | *Df* | *p value* |
| | | *LowC* | -434.54 | 6.5663 | 1 | **0.010** |
| | | *Standard* | -412.39 | 0.0841 | 1 | 0.772 |

| | | | | |
|---|---|---|---|---|
| *HighC* | -451.13 | 13.366 | 1 | **<0.001** |

**Table S2**. Summary of $log_{10}$ transformed bacterial load data of virgin and mated flies after 20 OD (equivalent to ~250 bacterial cells/fly) *P. rettgeri* infection, analyzed using a generalized linear model (GLM) using *Gamma distribution* (due to non-normal distribution of our data tested using Shapiro test) by fitting 'temperature' and 'dietary treatment' as categorical fixed-effects for both virgin and mated flies (Model: Log Avg *CFU~ Temperature× Dietary treatment, family=Gamma*). Statistically significant p-values have been highlighted in bold.

| | | ***Virgin*** | | | | ***Mated*** | | | |
|---|---|---|---|---|---|---|---|---|---|
| | ***Tested effect*** | *Chisq* | *Df* | *p value* | | *Chisq* | *Df* | *p value* | |
| *Full model* | *Temperature (T)* | 0.6327 | 1 | 0.729 | | 0.9223 | 1 | 0.337 | |
| | *Dietary treatment (DT)* | 0.0685 | 2 | 0.794 | | 3.1872 | 2 | 0.203 | |
| | *T ⨉ DT* | 0.0769 | 2 | 0.962 | | 0.1990 | 2 | 0.905 | |
| *Post hoc to compare dietary treatments across Temperatures* | *T* | *Estimate* | *SE* | *Z ratio* | *p-value* | *Estimate* | *SE* | *Z ratio* | *p-value* |
| | *Ambient* | | | | | | | | |
| | *LowC - HighC* | 0.1168 | 0.166 | 0.702 | 0.762 | 0.131 | 0.153 | 0.861 | 0.665 |
| | *LowC - Standard* | -0.0882 | 0.167 | -0.527 | 0.858 | -0.1212 | 0.153 | -0.792 | 0.708 |
| | *HighC - Standard* | 0.0286 | 0.163 | 0.176 | 0.983 | 0.0101 | 0.148 | 0.069 | 0.997 |
| | *Warm* | | | | | | | | |
| | *LowC - HighC* | 0.0679 | 0.163 | 0.416 | 0.909 | 0.225 | 0.147 | 1.532 | 0.276 |
| | *LowC - Standard* | -0.0263 | 0.165 | -0.159 | 0.986 | -0.161 | 0.149 | -1.080 | 0.526 |
| | *HighC - Standard* | 0.0416 | 0.162 | 0.257 | 0.964 | 0.064 | 0.141 | 0.455 | 0.892 |
| Post hoc to compare temperatures across dietary treatments | *DT* | *Estimate* | *SE* | *Z ratio* | *p-value* | *Estimate* | *SE* | *Z ratio* | *p-value* |
| | *LowC* | -0.0626 | 0.168 | -0.372 | 0.710 | -0.0349 | 0.156 | -0.224 | 0.823 |
| | *Standard* | -0.0007 | 0.164 | -0.004 | 0.996 | -0.0746 | 0.16 | -0.512 | 0.608 |
| | *HighC* | -0.0137 | 0.161 | -0.085 | 0.932 | -0.128 | 0.143 | -0.898 | 0.369 |

**Table S3**. Summary of the Cox proportional hazard model, fitting the model to estimate the response to starvation stress in virgin flies from an outbred population. The model is specified as survival ~ temperature x dietary treatment, with 'temperature' and 'dietary treatment' as fixed effects. The table shows model output (ANOVA) for survival against starvation in virgin flies. Statistically significant p-values are highlighted in bold.

| | ***Tested effect*** | *loglik* | *Chisq* | *Df* | *p value* |
|---|---|---|---|---|---|
| *Full model* | *Temperature (T)* | -2457.3 | 97.210 | 1 | **<0.001** |
| | *Dietary treatment (DT)* | -2450.9 | 12.991 | 2 | **0.002** |
| | *T × DT* | -2450.8 | 0.112 | 2 | 0.946 |
| *Post hoc to compare dietary treatments across Temperatures* | *T* | *loglik* | *Chisq* | *Df* | *p value* |
| | *Ambient* | | | | |
| | *LowC - HighC* | -657.36 | 6.4169 | 1 | **0.011** |
| | *LowC - Standard* | -669.78 | 1.9377 | 1 | 0.164 |
| | *HighC - Standard* | -665.31 | 0.6795 | 1 | 0.410 |
| | *Warm* | | | | |
| | *LowC - HighC* | -662.64 | 6.033 | 1 | **0.015** |
| | *LowC - Standard* | -648.82 | 3.1787 | 1 | 0.075 |
| | *HighC - Standard* | -650.12 | 0.5832 | 1 | 0.445 |
| Post hoc to compare temperatures across dietary treatments | *DT* | *loglik* | *Chisq* | *Df* | *p value* |
| | *LowC* | -650.13 | 31.042 | 1 | **<0.001** |
| | *Standard* | -640.61 | 29.76 | 1 | **<0.001** |
| | *HighC* | -642.13 | 36.88 | 1 | **<0.001** |

**Table S4**. Measurement of fertility for the flies from the outbred population following 24 hours of egg laying. Fertility was quantified by counting the number of offspring produced by 35 single females from each treatment. We analyzed the fertility using a generalized linear model (GLM) using a *Negative Binomial distribution* (due to the non-normal distribution of our data, tested using the Shapiro test) for the flies. We specified the model as: fertility ~ temperature x dietary treatment, with 'temperature' and 'dietary treatment' as fixed effects. Statistically significant p-values have been highlighted in bold.

| | ***Tested effect*** | *Chisq* | *Df* | *p value* | |
|---|---|---|---|---|---|
| *Full model* | *Temperature (T)* | 1.3352 | 1 | 0.248 | |
| | *Dietary treatment (DT)* | 13.8149 | 2 | **0.001** | |
| | *T × DT* | 5.8443 | 2 | 0.054 | |
| *Post hoc to compare dietary treatments across temperature* | *T* | *Estimate* | *SE* | *Z ratio* | *p-value* |
| | *Ambient* | | | | |
| | *LowC - HighC* | -0.6592 | 0.122 | -5.411 | **<0.001** |
| | *LowC - Standard* | 0.0199 | 0.116 | 0.172 | 0.984 |
| | *HighC - Standard* | -0.6393 | 0.122 | -5.242 | **<0.001** |
| | *Warm* | | | | |
| | *LowC - HighC* | -0.3081 | 0.067 | -4.607 | **<0.001** |
| | *LowC - Standard* | 0.2902 | 0.067 | 4.362 | **<0.001** |
| | *HighC - Standard* | -0.0179 | 0.0715 | -0.250 | 0.9661 |
| Post hoc to compare temperatures across dietary treatments | *DT* | *Estimate* | *SE* | *Z ratio* | *p-value* |
| | *LowC* | 0.143 | 0203 | 0.704 | 0.4816 |
| | *Standard* | 0.413 | 0.18 | 2.297 | **0.022** |
| | *HighC* | -0.208 | 0.155 | -1.349 | 0.177 |
